# Patterns of Structural Disconnection Driving Proprioceptive Deficits After Stroke

**DOI:** 10.1101/2025.05.22.655663

**Authors:** Mika Kaeja, Leila Gajiyeva, Yasser Iturria-Medina, Arno Villringer, Bernhard Sehm, Christopher Steele

## Abstract

**Background:** Stroke is a leading cause of death and disability worldwide, with proprioceptive impairments affecting up to 64% of survivors. These impairments hinder sensorimotor function and motor recovery, significantly impacting post-stroke disability and quality of life. Proprioception depends on an integrated network of brain regions but remains underexplored due to limitations in clinical assessments, making it difficult to identify precise links between stroke-related damage and functional deficits. To address this, we combined quantitative proprioceptive measurements with Connectome-based Lesion-Symptom Mapping (CLSM) to identify white matter (WM) disconnection patterns underlying proprioceptive deficits following sensorimotor stroke.

**Methods:** In this observational study, we investigated the relationship between WM disconnection and proprioceptive deficits in 39 chronic stroke survivors with paretic arm function (>6 months post-stroke; 13 females; aged 35-81) using CLSM and kinematic assessments. Lesions were manually delineated from 3T MRI scans, and proprioception was quantified using the Arm Position Matching (APM) task on the KINARM Exoskeleton. Patient-specific voxel-wise WM disconnection maps were generated using the Tractography Lesion Assessment Standard (TractLAS), which quantifies disconnection relative to a healthy WM connectome. Proprioceptive scores were regressed against disconnection maps using voxel-wise linear regressions (family-wise error corrected, controlled for age and sex).

**Results:** Our disconnectome-based approach identified a network of regions where proprioceptive deficits were significantly associated with WM disconnection (d = 0.55-1, p < .005 FWE, t = 3.48-6.35). These included tracts previously implicated in proprioceptive function (superior longitudinal fasciculus, middle longitudinal fasciculus, and arcuate fasciculus) and beyond (medial lemniscus, spinothalamic tract, posterior thalamic radiation).

**Conclusion:** We provide evidence that post-stroke proprioceptive impairments arise from network-wide WM disconnection in several key tracts that mediate proprioceptive function. This study highlights the benefits of using CLSM to assess stroke-related proprioceptive deficits and offers a framework for network-informed assessments of functional impairments that can be used for targeted therapies post-stroke.

## Introduction

Stroke is one of the leading causes of death and disability across the globe,^1^ with survivors experiencing a wide range of impairments including sensory, motor and cognitive deficits, which are dependent on the size and location of lesions.^2–4^ Proprioception is one crucial but underexplored function that is frequently affected by stroke, significantly impacting motor recovery and everyday life of patients. Indeed, up to 64% of stroke survivors are affected by proprioceptive deficits,^5^ which is concerning as proprioception serves a key role in re-learning motor skills and improving functional outcomes.^6–9^ However, proprioception is difficult to accurately assess with conventional clinical tests,^7,10^ leaving the impact of stroke-induced White Matter (WM) disconnection on proprioceptive deficits completely unexplored. Given that proprioceptive function requires the integration of information across multiple brain regions,^11^ a more holistic connectome-informed understanding of stroke-related deficits is crucial for tailoring interventions to facilitate recovery.

Stroke-induced proprioceptive deficits not only affect motor coordination but also impede the ability to learn or re-learn the motor skills necessary to improve functional outcomes, negatively impacting a patient’s recovery trajectory, personal autonomy, and quality of life.^12^ Despite the importance of proprioceptive abilities in stroke motor recovery, standard clinical tools use ordinal scales with non-independent measures (i.e the Nottingham Sensory Assessment)^13^ that are not easily quantifiable and therefore hamper direct comparisons between/across patients.^7,10^ Kinematic assessments have been proposed to solve these drawbacks and can be used to accurately and quantitatively assess proprioceptive deficits and the evolution of recovery.^8,10,14^ The Arm Position Matching Task (APM),^7^ conducted using the KINARM robot, is a validated tool for quantifying proprioceptive deficits in stroke patients.^14–16^ Using quantitative assessment allows for a deeper and more precise understanding of the neural underpinnings of proprioception.

Proprioception relies on the integration of information across a network of brain regions - including the supramarginal gyrus (SMG), superior temporal gyrus, lateral thalamus and somatosensory cortex – which are thought to support the perception of limb position.^11,17,18^ These regions are connected by key white matter tracts such as the superior longitudinal fasciculus (SLF), middle longitudinal fasciculus (MdLF), arcuate fasciculus (AF), and the dorsal column of the medial lemniscus.^16,19^ Stroke-induced damage can disrupt both gray matter regions and the underlying WM network, leading to functional impairments.^18^ A connectome-based approach, which examines how local stroke damage affects the WM network beyond the lesion boundaries (i.e., connectome-based lesion symptom mapping [CLSM]), is therefore a valuable approach for understanding the global effects of damage^2,20^ and is particularly relevant for analyzing proprioception due to its integrative function across the brain.

In this study, we identify how damage to specific regions of the WM network is related to quantified deficits in proprioceptive function using CLSM (TractLAS)^21^ and kinematic measurements^7,10^ in chronic sensorimotor stroke survivors. Further, our approach provides a holistic network-informed assessment of quantified impairments that can identify the underlying WM network affected in individual patients and have the potential to be used for the design of patient-specific rehabilitation.

## Method

### Clinical Population

#### Participants

Forty-two (16 females) first-time sensorimotor ischemic stroke patients in the chronic stage were recruited from the University Hospital Leipzig Day Clinic for Cognitive Neurology (UHL-DCCN) in Leipzig, Germany (aged 35-81, M = 61 ± 11 yo). Inclusion criteria were having 1) their first sensorimotor stroke at least 6 months previously and 2) a Fugl-Meyer Upper Limb Assessment between 20-58 points (detailed in Supplementary Table S1). Exclusion criteria were 1) patient was unable to follow instructions due to neurological disease, 2) a Mini-Mental State Examination score under 24 or experiencing visual impairments and 3) contraindication to Magnetic Resonance Imaging (MRI). The ethics certification for this study was granted by the University of Leipzig’s Medical Faculty Ethics Committee and participants provided written informed consent according to the Declaration of Helsinki.

#### Apparatus

Proprioception was measured using the KINARM Exoskeleton Lab (BKIN Technologies Ltd., Kingston, ON, Canada) which records the pattern of joint movements and muscular torques produced by the upper extremities. Both upper limbs were fully supported by the robot at shoulder level while patients moved within an integrated 2D augmented reality display, allowing for presentation and interaction with visual targets in the same plane as limb motion. Task data was collected with Dexterit-E™ v 3.5.3 and analysed with v 3.6.2.

#### Proprioceptive Assessment

The Arm Position Matching Task (APM)^7^ from the KINARM Standard Task Library (BKIN Technologies Ltd., Kingston, ON, Canada) was used to assess proprioceptive impairment. While both arms were obstructed from view, the patient was asked to relax their affected hand and allow the robot to move it to one of nine randomized locations in a 3x3 square (Figure 1). Participants were instructed to mirror match that location in space with their unaffected hand and to notify the experimenter when they judged that they had reached the mirrored position. Patients underwent a total of 54 APM trials over the course of a single session. The summary task score was used to operationalize patient’s performance, where greater scores represented greater proprioceptive impairment.

**Figure 1.**
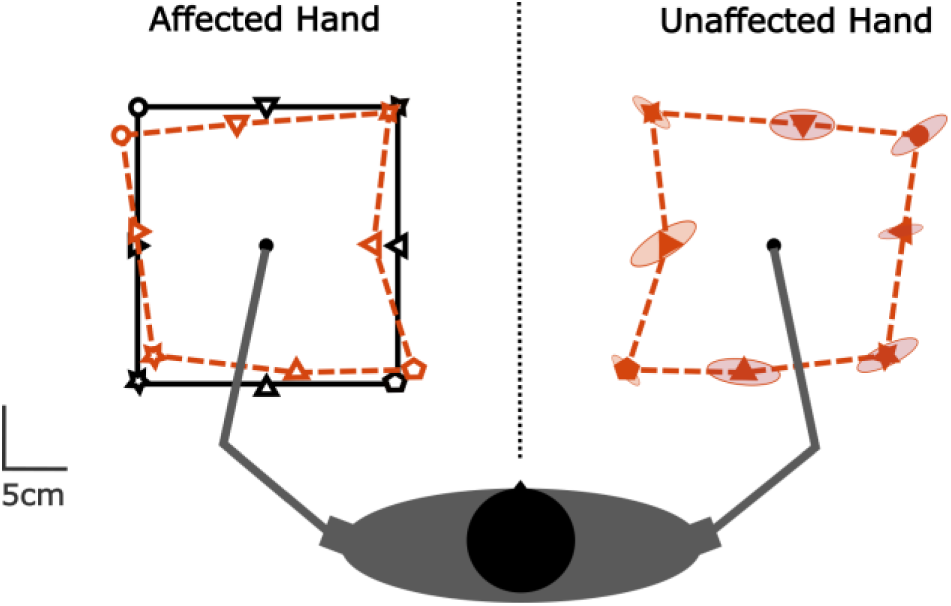
Arm Position Matching Task. This task assesses proprioception as patients use their unaffected hand (right) to mirror match the position of their affected hand (left) after it has been moved by a robot to one of nine spatial locations (black symbols). The solid orange symbols on the right represent examples of the participant’s attempted matches. Accuracy in this task reflects proprioceptive function and helps assess their degree of impairment (Adapted from BKIN Technologies Ltd., 2021).

#### MRI Acquisition

Chronic stroke patients from the UHL-DCCN were briefed about the study procedures and intent and provided written informed consent to participate. Following the acquisition of consent, MRI scans were acquired using a 3T Siemens MRI with a 32-channel head coil (Siemens Healthineers AG, Forchheim, Germany). Acquisition sequences included T2 Fluid Attenuated Inversion Recovery Imaging (FLAIR; anisotropic resolution = 1 x 0.49 x 0.49 mm, echo time = 395), Diffusion Weighted Imaging (DWI; anisotropic resolution = 1.72 x 1.72 x 1.7 mm) and T1-weighted imaging (T1w; isotropic resolution = 1 mm, repetition time = 4.21).

#### MRI Processing

Lesions were segmented by hand on the FLAIR images. Segmentations were initially performed by a doctor in training (L.G.), individually verified by a clinical neurologist (B.S.), and then binarized to form the final lesion mask for each patient. Lesions in the left hemisphere were flipped into the right for subsequent analysis. The lesion overlap between patients in Montreal Neurological Institute (MNI) space, visualised using MRtrix3,^22^ is shown in Figure 2. T1w images were linearly registered to the standardized MNI space using Advanced Normalization Tools (ANTs).^23^ Lesion masks were normalized to the final connectome model space by concatenating the appropriate transforms (nearest neighbour interpolation).

**Figure 2.**
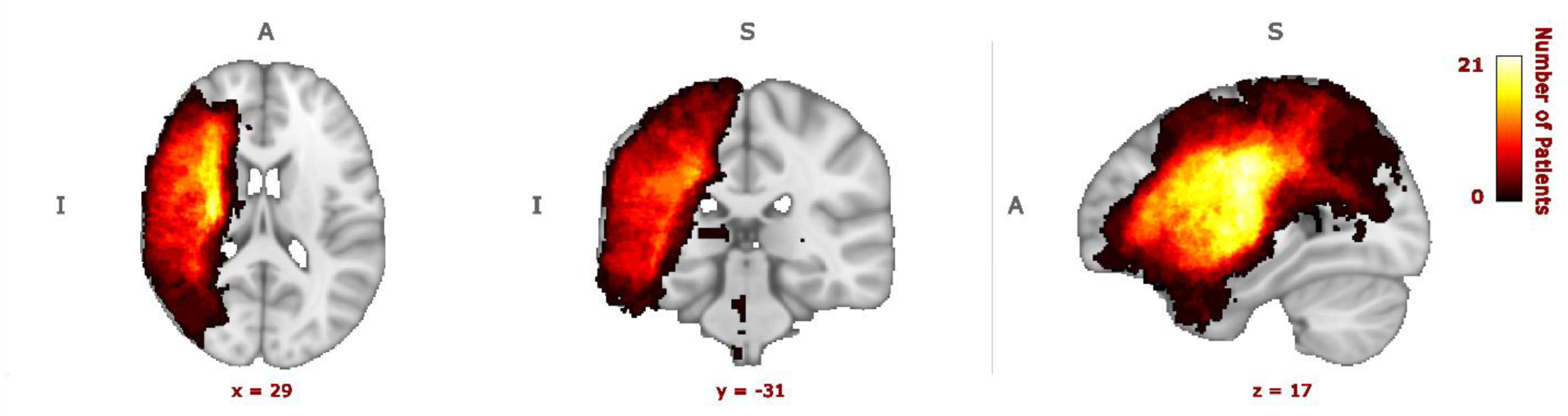
**Lesion Overlap Across All Patients**. Lesion overlap across all 39 patients with APM values. The area of maximal overlap is across the sensorimotor system. All lesions were flipped into the right hemisphere for analyses. The colourbar indicates lesion overlap from 1 (dark red) to 21 (bright yellow) patients. Axis labels are as follows: A = anterior, P = posterior, S = superior, I = ipsilesional hemisphere.

#### Connectome Model

To allow direct comparisons across individual patients we generated a biologically plausible connectome model to serve as a quantified baseline for comparison, as has previously been proposed in our^21^ and others’ work.^20,24,25^ The connectome model was based on the data of 1001 participants (aged 22-37, M = 29 ± 4 yo) from the Human Connectome Project – Young Adult (HCP-YA; HCP S1200 release).^26^ The imaging acquisition procedures of the HCP-YA data can be found in Van Essen et al,^26^ while the preprocessing steps used to construct the model using MRtrix3^22^ are fully described in Tremblay et al.^27^ In short, T1w images were segmented into five tissue types (5tt) and the minimally preprocessed DWI data was processed with Constrained Spherical Deconvolution analysis (CSD)^28^ to generate Fibre Orientation Distribution functions (FODs). FODs were used in conjunction with the 5tt segmentation to perform group registration that optimized for the alignment of FODs within white matter. The final connectome model was computed by performing anatomically constrained tractography^29^ on the FOD optimized group average (detailed in Supplementary Methods). To maximize biological plausibility,^30,31^ we first generated a very large number of oversampled streamlines that initiated and terminated in the WM-GM interface (100 million). We then performed two sets of filtering: first selecting the 10 million most plausible streamlines^30^ and then reweighting them^31^ to fit the original FODs in each voxel. This process works globally to ensure that the resulting streamlines (and their associated weights) are constrained by the underlying diffusion data and are, therefore, more biologically plausible.^31^

#### Patient Specific Disconnectomes

Each patient’s lesion mask was introduced into the connectome model as an exclusion mask to remove any streamlines that would have passed through lesioned tissue. The affected streamline counts were divided by the density of connections at each voxel relative to a healthy connectome model to calculate each patient’s proportion of stroke-induced disconnection in every voxel of the brain (i.e., their disconnectome).^21^. As the disconnectome is computed relative to a healthy connectome model, this method can be used to compare across patients regardless of the specific lesion locations. Our approach, the Tractography-based Lesion Assessment Standard (TractLAS),^21^ is composed of a custom-built Python script that calls specific MRtrix3 functions. Figure 3 provides a visual summary of this approach, while further methodological details and patient-specific results are available in Supplementary Methods and Figure S1.

**Figure 3.**
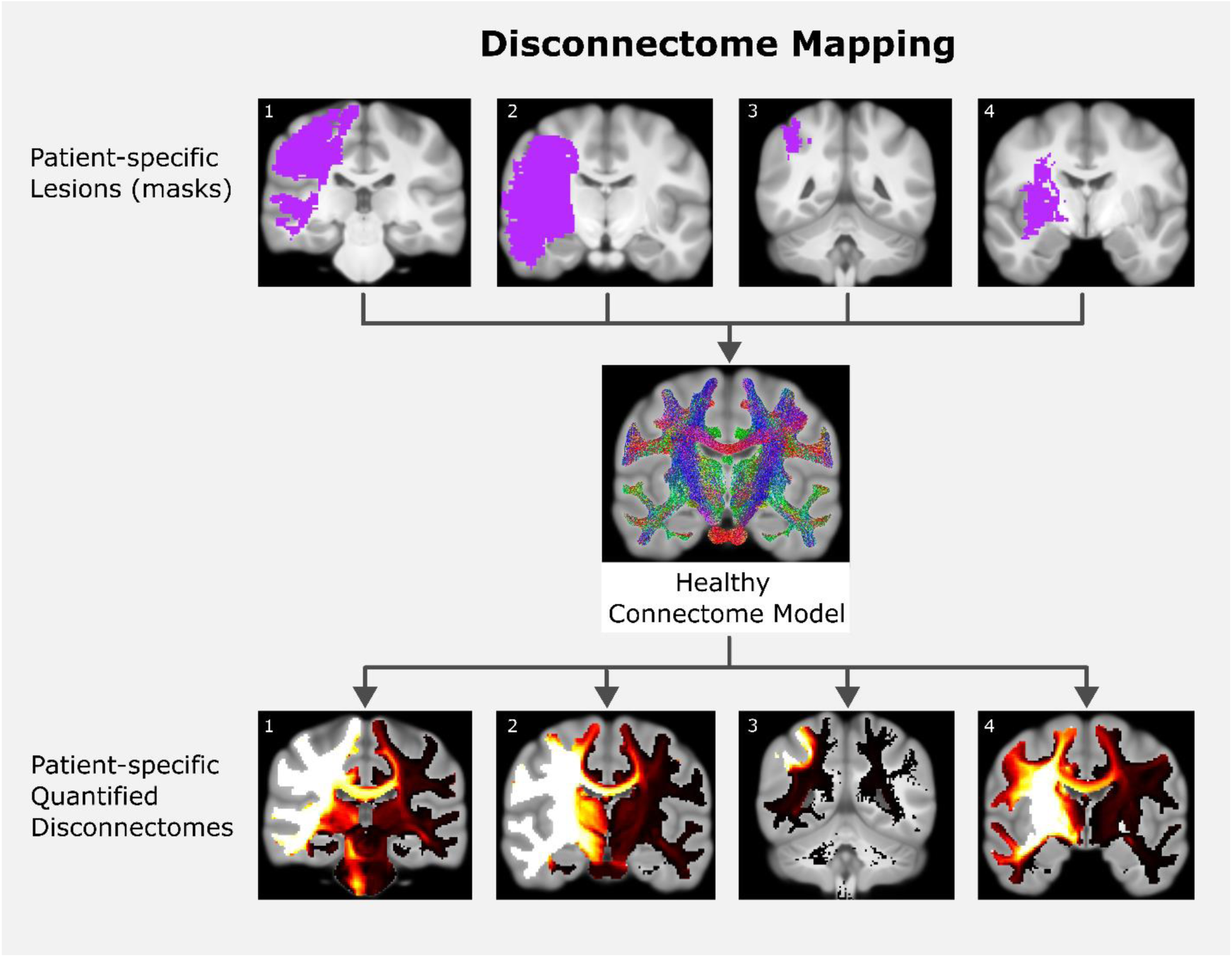
A visual summary of the disconnectome mapping approach with patient-specific examples. The lesion masks of each patient (seen in purple) are individually projected into the healthy connectome model to obtain each of their quantified disconnectome’s (maximum disconnection proportion in yellow-white). The corresponding numbers on the lesion masks disconnectomes indicate the same patients. Please refer to Supplementary Figure S1 for all patient disconnectomes.

### Statistical Analysis

To identify WM tracts where disconnection is associated with proprioceptive deficits, we performed mass univariate voxel-wise regressions in which APM scores of the affected limb served as the dependent variable and the proportion of disconnection at each voxel as the independent variable. Three patients did not complete the APM due to fatigue and were excluded at analysis, yielding a final sample of 39 with complete data for both variables. These variables were entered into ordinary least-squares general linear regressions with Threshold Free Cluster Enhancement (TFCE)^32^ and 5,000 randomized permutations using Nilearn^33^ with a rigorous family-wise error (FWE) correction threshold of p<0.005 while controlling for age and sex.

Regional identifications of significant results were made with reference to the Johns Hopkins University Diffusion International Consortium of Brain Mapping WM parcellation atlas (JHU-ICBM)^34^, the Harvard-Oxford cortical and subcortical structural atlases,^35^ and the Brainstem Connectome Atlas.^36^ In addition, T-values were converted to Cohen’s d values to calculate effect size, and regions with moderately strong effects (d > 0.6) were subsequently presented for display using MRTrix3^22^ and Nilearn^33^ Additional details for the specific functions used can be found in the Supplementary Methods section.

## Results

Thirty-nine out of 42 chronic sensorimotor stroke patients (13 females) aged 35-81 (*M* = 59.95 ± 10.86) successfully completed the APM task and exhibited a wide range of performance (range: 0.01-5.39, *M* = 1.83 ± 1.29; refer to Supplementary Table S1 for details). As expected, lesion overlap across patients in our sensorimotor stroke group was largest in white matter underlying the sensory, motor, and premotor cortices (Figure 2). Each patient-specific disconnectome exhibited distinct disconnection patterns, which can be seen in Supplementary Figure S1.

The analysis regressing proprioceptive impairment and disconnection values, while accounting for age and sex, revealed an extensive set of significant WM regions (d = 0.55-1, p < .005 FWE, t = 3.48-6.35) underlying the parietal, supplementary motor, temporal, precentral, postcentral and supramarginal cortices (Figure 4 & 5). To focus on the key white matter regions supporting proprioception, we present a subset of the significant areas (along with their general function) that exceeded a moderate effect size (d > 0.6) in Table 1. This cutoff corresponds to a FWE corrected p < .003 and t > 3.80 as shown in Figure 4 and 5.

**Figure 4.**
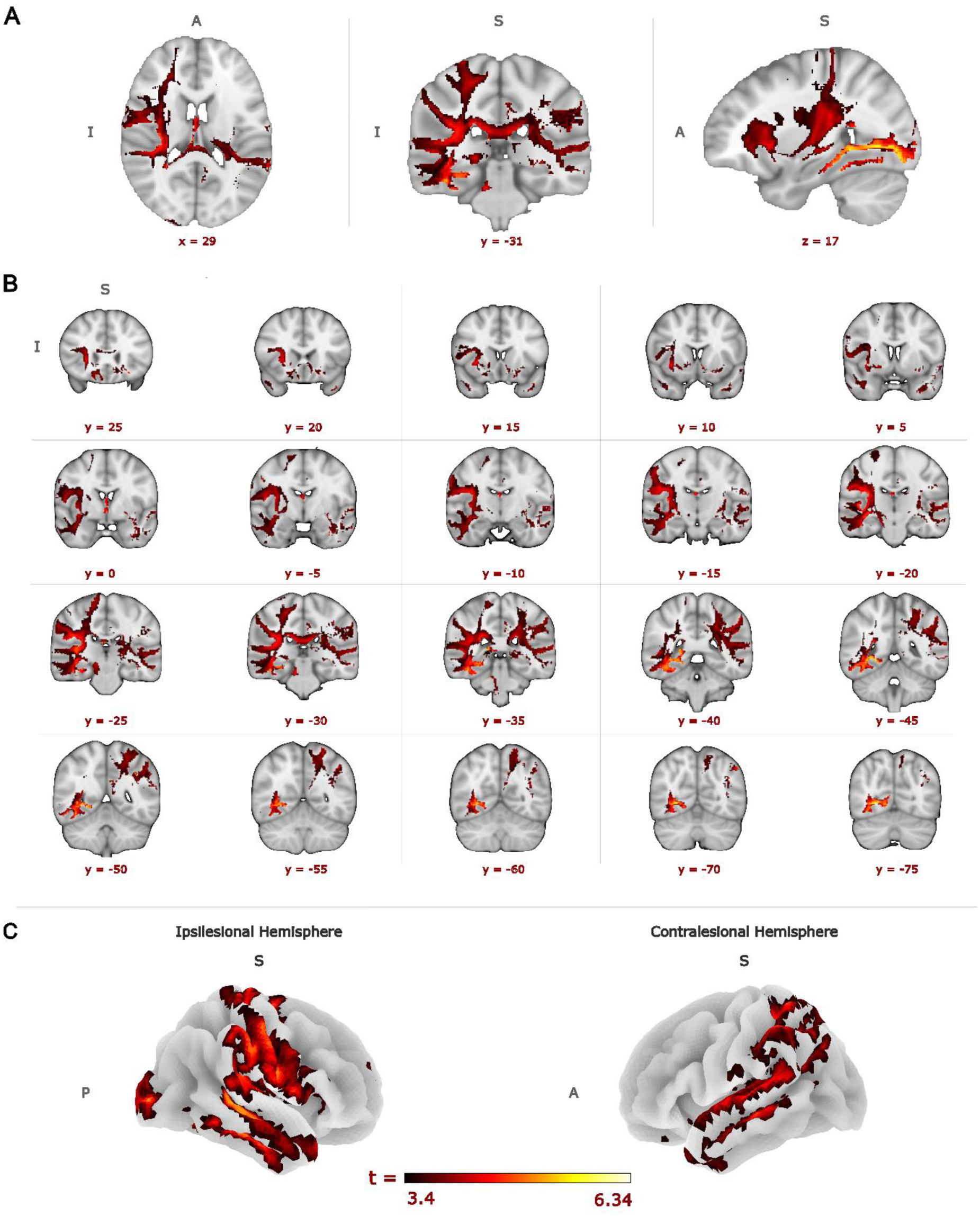
Relationship between stroke-induced disconnection and proprioceptive impairment. Significant voxel-wise t values (p < .005, t > 3.48) showing associations between proprioceptive impairment and stroke-induced WM disconnection. (A) Orthogonal views and (B) corresponding coronal slices of the proprioceptive disconnection network. All images are in radial convention in MNI space (mm), with the affected hemisphere on the radiological right (R). (C) Projected cortical surface effects of the proprioceptive disconnection network. Darker colours indicate lower t values; lighter colours indicate higher t values. Axis labels are as follows: A = anterior, P = posterior, S = superior, I = ipsilesional hemisphere.

**Figure 5.**
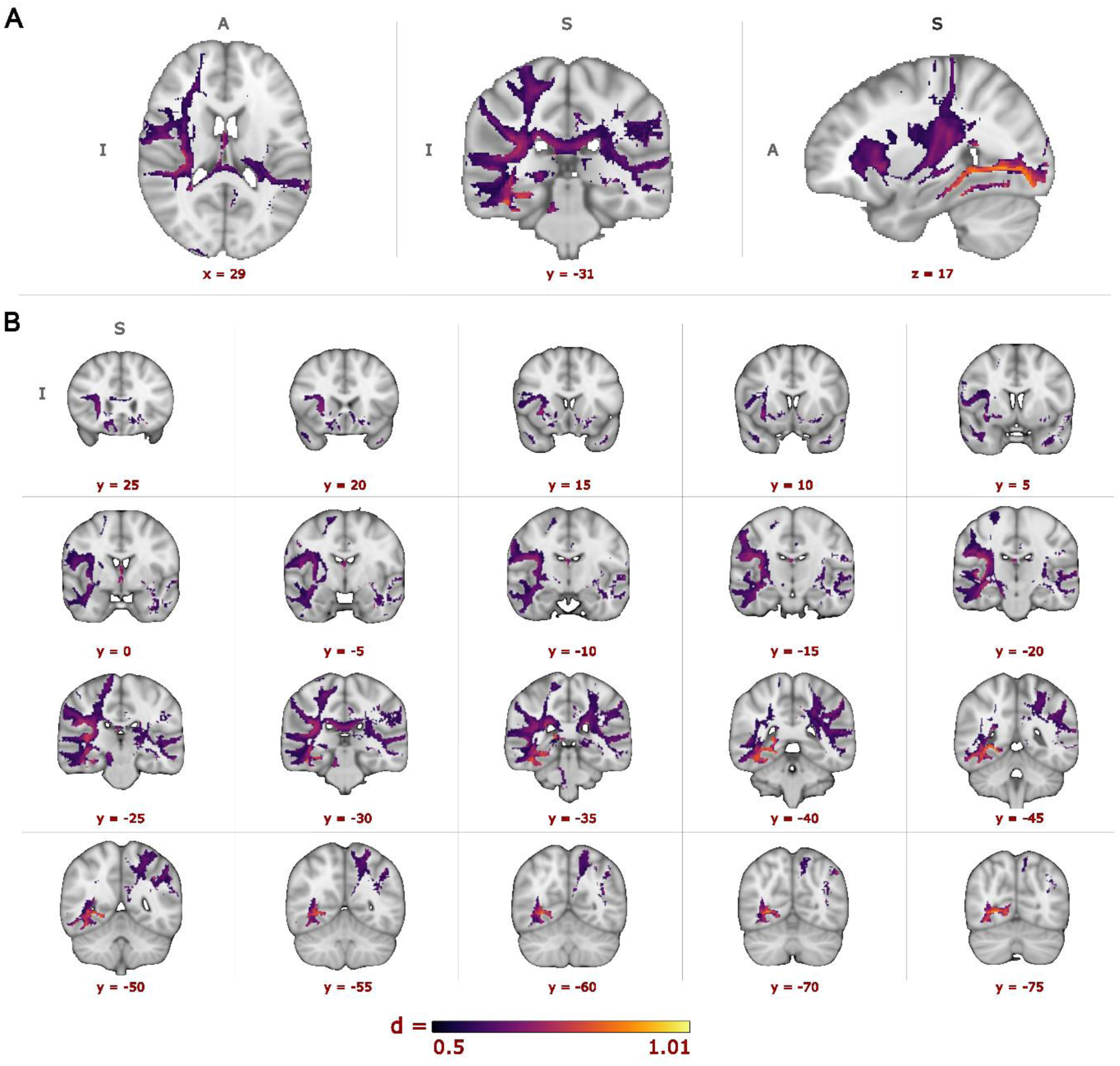
Effect size associated with the relationship between stroke-induced disconnection and proprioceptive impairment. Cohen’s d values (p < .005 d > .55) showing the voxel-wise effect size associated with the relationship between proprioceptive impairment and stroke-induced disconnection. (A) Orthogonal views and (B) corresponding coronal slices of the proprioceptive disconnection network. Darker colours indicate lower Cohen’s d values; lighter colours represent higher Cohens d values. All images are in radial convention in MNI space (mm), with the affected hemisphere on the radiological right (R). Acronym labels are as follows: A = anterior, P = posterior, S = superior, I = ipsilesional hemisphere.

**Table 1.**
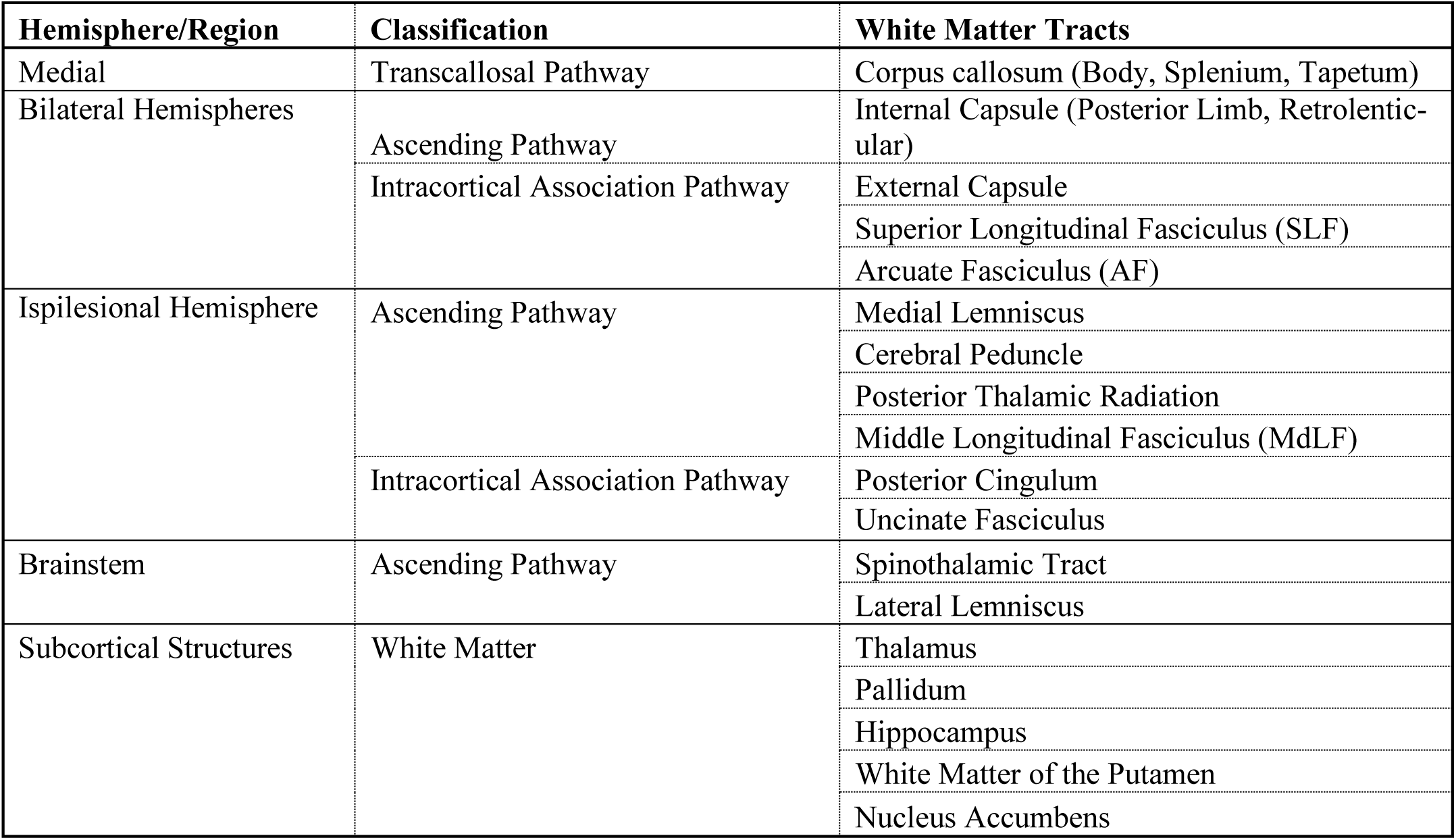
White matter tracts supporting proprioception post-stroke. . List of white matter tracts impacted by stroke-induced disconnection and impaired proprioceptive function post- stroke, significant at the d > 0.6, p < 0.003, t > 3.80 level.

## Discussion

Our findings reveal, for the first time, the brain-wide structural white matter disconnection patterns underlying proprioceptive deficits after sensorimotor stroke. By mapping patient-specific disconnection profiles, we provide evidence that proprioceptive impairments arise from disruptions across a distributed white matter network, rather than a single localized region. Given that proprioception requires integration across multiple brain regions,^11^ our results highlight the importance of a whole-brain connectivity approach in assessment. This framework not only deepens our understanding of stroke-related proprioceptive deficits but also aids in identifying potential intervention targets. Ultimately, these findings underscore the importance of precise behavioral quantification and emphasize the need to consider white matter network disconnection when assessing proprioceptive impairments and designing rehabilitation strategies post-stroke.

### Network-Wide White Matter Disruptions in Post-Stroke Proprioception

We identified an expanded network of white matter regions involved in post-stroke proprioceptive function, reaffirming the role of previously implicated pathways while highlighting additional contributors.^16,18,37,38^ Specifically, both our findings and prior research have identified the SLF, AF and MdLF as key pathways associated with proprioceptive function. Chilvers et al. (2022) found that lower fractional anisotropy in these tracts, as assessed using the APM task, was associated with poorer proprioceptive performance – a finding supported by our results, which showed strong associations between damage to these pathways and proprioceptive impairments. This likely reflects disconnection within white matter pathways responsible for integrating key cortical regions involved in proprioceptive awareness of limb location: the SLF links the superior parietal lobule with premotor and primary motor areas (M1); the AF connects posterior temporal regions with the prefrontal cortex; the and MdLF interconnects the superior temporal gyrus and parietal regions, supporting multimodal sensory integration.^16,18,37^ These salient tracts extend well beyond the boundaries of patients’ lesions, suggesting that proprioceptive deficits arise from network-wide white matter disconnection induced by sensorimotor stroke.

Across the global network of the brain, many tracts must function in tandem to effectively perform proprioceptive tasks.^11^ Based on our findings, several ascending, transcallosal and intracortical WM tracts appear to be implicated in post-stroke proprioceptive processing and warrant further exploration. First, we found that the medial lemniscus and spinothalamic tracts, both key ascending pathways known to transmit proprioceptive and somatosensory information, were significantly associated with proprioceptive scores in our stroke cohort. Although these tracts have been theoretically linked to proprioception,^39^ our results provide the first direct evidence of their involvement in post-stroke proprioceptive deficits. In addition, our analysis revealed significant associations with ascending pathways, posterior limb and retrolenticular part of the internal capsule, cerebral peduncle, anterior and posterior thalamic radiation; transcallosal tracts including the body, splenium and tapetum of the corpus callosum; intracortical association pathways such as the external capsule, posterior cingulum, uncinate fasciculus; and WM within the nucleus accumbent, thalamus, pallidum, putamen, and hippocampus. These regions, many of which are known to support sensorimotor integration, interhemispheric communication, and related cognitive and limbic processes, likely contribute to the integrity of the post-stroke proprioceptive network.^40–45^ Notably, while the ipsilesional hemisphere showed more focal associations with classical ascending sensory pathways, the contralesional hemisphere contributed through widespread transcallosal and intracortical association pathways, suggesting a potential compensatory role in network-level reorganization. Together, our results strongly suggest that post-stroke proprioceptive function relies on a distributed network of ascending, transcallosal and intracortical white matter pathways, several of which have not been previously recognized in this context.

### Towards Patient-Centred Rehabilitation

To fully understand the impact of lesions on proprioception, it will be important to consider both individual patient-specific disconnectomes and group-level results. The patient-specific disconnectomes allow us to identify the precise white matter pathway(s) disrupted in each patient (therefore highlighting the unique damage profile that is related to their potentially complex behavioural deficits), while group analyses provide key results for identifying core regions that can be generalized. Since lesions vary greatly in size and location, incorporating patient-specific disconnection patterns into clinical frameworks could inform more tailored rehabilitation approaches. This combined methodology holds promise for identifying individualized intervention targets and moving toward more nuanced, patient-centered care. As rehabilitation plays a critical role in post-stroke recovery and long-term quality of life, disconnectome-informed intervention planning may offer a valuable avenue for improving functional outcomes.^46^

## Limitations, Implications, and Future Directions

One limitation of our approach is that disconnectomes are computed relative to a model that, like any model, is an approximation. Our model was built from high resolution diffusion MRI data of healthy young adults, and while this data is of very high quality it may therefore reflect both biases in the “normal” young adult brain and in the diffusion MRI data and tractography method that we have used. However, mitigating against this issue is that our approach computes all disconnectomes relative to the same model and therefore ensures that different patients are directly comparable and, if present, suffer from the same biases and/or inaccuracies.^47^

Beyond these methodological limitations, the generalisability of our findings may also be constrained by the study sample and imaging context. All participants were chronic ischemic stroke survivors with sensorimotor impairments, which may limit applicability to patients in acute or subacute stages, or those with hemorrhagic or cognitive-dominant strokes. Additionally, the high-resolution diffusion MRI data used to derive disconnectomes may not be widely available in routine clinical settings, potentially limiting real-world implementation.

Quantifying individual patient-specific disconnections that result from specific lesion locations can serve as a useful guide towards developing targeted rehabilitative approaches. Knowledge of each patient’s disconnectome, considered within the context of their deficits, can be used to develop a comprehensive neurorehabilitation regimen using physical (bimanual training)^48^, mental (cognitive training), medical (activity modulating drugs) and technologically assistive (brain-computer interface, deep brain stimulation, non-invasive brain stimulation, and robotic training)^49–51^ techniques that target white matter tracts (and their linked cortical regions) most affected in each patient. This regiment could strengthen their weakened connections and provide support to damaged compensatory pathways to allow them to efficiently recover a degree of function that has been lost. We can additionally look longitudinally at the association between the effectiveness of rehabilitation techniques and individual patterns of disconnection with their resulting impairments. This approach could be used to create a predictive model of the most effective rehabilitation techniques to create a personalized rehabilitation program based on their pattern of disconnection and associated behavioural deficits. In this way, commonly available imaging measurements (i.e., lesion masks) could give practitioners insight into predicted stroke outcomes and the most effective interventions early in recovery.^52^

## Conclusion

This work is the first to explore how stroke-induced WM disconnection affects proprioceptive impairments, identifying the key white matter networks supporting proprioception after sensorimotor stroke. Our novel approach allowed us to identify strong relationships with white matter disconnection consistent with the existing literature,^16,18,37,38^ and significantly extends this work by both highlighting the key importance of the underlying WM connecting regions and identifying putative additional tracts for further study. Our results provide strong evidence that behavioural deficits are differentially affected by the location of lesions and their resulting WM disconnections. Ultimately, the current research strongly supports the use of disconnectome mapping to accurately characterize individual differences in the behavioural effects of stroke lesions and allow for direct comparisons between patients with differing lesion locations.

## Acknowledgements

The authors would like to thank the patients at the Max Planck Institute for Human Cognitive Brain Science Neurology Clinic for allowing the use of their data for this study, and Dr. Bernhard Sehm for providing it. We also thank Zaki Alasmar for assistance with the pilot methodology of this project.

## Sources of Funding

This research was funded by operational grants received by Christopher J. Steele from the Natural Sciences and Engineering Research Council of Canada (DGECR-2020-00146), the Canadian Institutes of Health Research (HNC 170723), the Canada Foundation for Innovation (CFI-JELF Project number 43722) and Heart and Stroke Foundation of Canada (National New Investigator.

## Disclosures

The authors have no competing interests to declare.

## Data, Materials, and Code Disclosure

Statistical maps will be available from the corresponding author upon reasonable request and released on Neurovault following publication. Raw data cannot be made available due to the ethical standards of the Ethics Committee of the University of Leipzig.

## Supplementary Materials

Supplemental Methods, describing in full the methodological and analytical techniques used in this study; Supplemental Table S1, a table describing patients’ demographic information and behavioural scores; Supplemental Figure S1, Each patient’s lesion and their resulting TractLAS generated disconnectomes.

## Non-standard Abbreviations and Acronyms

5tt: Five Tissue Types
AF: Arcuate Fasciculus
ANTs: Advanced Normalization Tools
APM: Arm Position Matching Task
CLSM: Connectome-based Lesion Symptom Mapping
CSD: Constrained Spherical Deconvolution analysis
DWI: Diffusion Weighted Imaging
FLAIR T2: Fluid Attenuated Inversion Recovery Imaging
FODs: Fibre Orientation Distribution functions (FODs)
FWE: Family-wise Error
GLR: General Linear Regressions
GM: Grey Matter
HCP-YA: Human Connectome Project – Young Adult
JHU-ICBM: Johns Hopkins University Diffusion International Consortium of Brain Mapping
MdLF: Middle Longitudinal Fasciculus
MNI: Montreal Neurological Institute
MRI: Magnetic Resonance Imaging
SLF: Superior Longitudinal Fasciculus
T1w T1: weighted Imaging
TFCE: Threshold Free Cluster Enhancement
TractLAS: Tractography-based Lesion Assessment Standard
UHL: DCCN University Hospital Leipzig Day Clinic for Cognitive Neurology
WM: White Matter

## Notes

### Competing Interest Statement

The authors have declared no competing interest.

